# Study of mitotic chromatin supports a model of bookmarking by histone modifications and reveals nucleosome deposition patterns

**DOI:** 10.1101/233056

**Authors:** Elisheva Javasky, Inbal Shamir, Shashi Gandhi, Shawn Egri, Oded Sandler, Scott B. Rothbart, Noam Kaplan, Jacob D. Jaffe, Alon Goren, Itamar Simon

## Abstract

Mitosis encompasses key molecular changes including chromatin condensation, nuclear envelope breakdown, and reduced transcription levels. Immediately after mitosis, the interphase chromatin structure is reestablished and transcription resumes. The reestablishment of the interphase chromatin is probably achieved by ‘bookmarking’, *i.e.*, the retention of at least partial information during mitosis. Yet, while recent studies demonstrate that chromatin accessibility is generally preserved during mitosis and is only locally modulated, the exact details of the bookmarking process and its components are still unclear. To gain a deeper understanding of the mitotic bookmarking process, we merged proteomics, immunofluorescence, and ChIP-seq approaches to study the mitotic and interphase genomic organization of human cells. We focused on key histone modifications and employed HeLa-S3 cells as a model system. Generally, we observed a global concordance between the genomic organization of histone modifications in interphase and mitosis, yet the abundance of the two types of modifications we investigated was different. Whereas histone methylation patterns remain highly similar, histone acetylation patterns show a general reduction while maintaining their genomic organization. These results demonstrate that the epigenomic landscape can serve as a major component of the mitotic bookmarking process. Next, to further investigate mitosis-associated chromatin changes, we followed up on previous studies that showed that nucleosome depleted regions (NDRs) become occupied by a nucleosome during mitosis. Surprisingly, we observed that the nucleosome introduced into the NDR during mitosis encompasses a distinctive set of histone modifications, differentiating it from the surrounding nucleosomes. We show that the nucleosomes near the NDR appear to both shift into the NDR during mitosis and adopt a unique modification pattern. HDAC inhibition by the small molecule TSA reverts this pattern. These results provide evidence for a mitotic deposition and change in the modifications of the nucleosomes surrounding the NDR. Altogether, by merging multiple approaches, our study provides evidence to support a model where mitotic bookmarking is achieved by histone modifications and uncovers new insights into the deposition of nucleosomes during mitosis.

## Introduction

The cell cycle is a dynamic orchestrated process that involves major changes to the chromatin structure (Raynaud et al. 2014) and the expression of hundreds of genes (Spellman et al. 1998; Cho et al. 2001; Whitfield et al. 2002; Bar-Joseph et al. 2008). During the cell cycle, DNA is replicated along S-phase and then condensed in mitosis prior to cell division.

During mitosis the cell undergoes profound changes. The nuclear envelope breaks apart and the chromosomes change their shape into distinguishable thick structures. Multiple studies have examined the structure of mitotic chromosomes using microscopy (Vagnarelli 2013), but they lack fine-detail information on individual loci. Recently, genome-wide long-range interactions within mitotic chromosomes were investigated by HiC and 4C (Naumova et al. 2013; Dileep et al. 2015; Stevens et al. 2017). These studies demonstrate the disappearance of the consistent higher-order chromatin structure in mitosis, including both genome compartments and topological associated domains (TADs). Nevertheless, the unique features of mitotic chromatin at a finer genomic resolution remain unexplored. In addition to chromosome reshaping, mitotic chromosomes are characterized by the disassociation of numerous proteins including RNA polymerase II and many gene-specific and general transcription factors (Martinez-Balbas et al. 1995; Parsons and Spencer 1997; Kadauke and Blobel 2013). The disassociation of these transcription machinery components leads to a dramatic decrease in transcription (King and Barnhisel 1967; Gottesfeld and Forbes 1997; Liang et al. 2015; Palozola et al. 2017).

Immediately after mitosis, the chromatin structure is gradually reestablished – the chromosomes decondense throughout G1 (Belmont and Bruce 1994), the interphase TAD structures are reformed within four hours after cytokinesis (Dileep et al. 2015), and transcription is reinitiated at the beginning of G1 (Prescott and Bender 1962; Gottesfeld and Forbes 1997; Prasanth et al. 2003). This complex orchestration of TAD re-formation and controlled transcription reinitiation necessitates precise epigenomic markings, which are likely preserved during mitosis despite the radical changes to the chromatin. How is this epigenomic information stored during mitosis? The current view is that partial information is retained on mitotic chromosomes to serve as bookmarks for the reestablishment of the chromatin structure. In line with this view, even in their condensed state, mitotic chromosomes remain accessible as is evident by both biophysical measurements (Chen et al. 2005; Caravaca et al. 2013) and DNase I sensitivity assays (Gazit et al. 1982; Kadauke et al. 2012). Furthermore, genomic chromatin accessibility maps by DNase I sensitivity and ATAC-seq comparing mitotic and interphase cells revealed that accessibility is generally preserved and is only locally modulated during mitosis (Hsiung et al. 2015; Teves et al. 2016). Yet, it remains unclear what serves as the bookmarks for the restoration of the global chromatin structure following the changes that occur during mitosis.

Much effort has been devoted to identifying components of the transcription machinery that may serve as cell cycle bookmarks. These studies focused mainly on general and specific transcription factors that are retained at a subset of their locations during mitosis (reviewed in (Kadauke and Blobel 2013; Zaidi et al. 2014)). Additionally, preliminary evidence suggests that histone tail modifications themselves participate in carrying out the task of bookmarking. It has been reported that retaining H4K5ac levels during mitosis is crucial for the bookmarking of a reporter gene (Zhao et al. 2011). Further, the global levels of several histone modifications were shown to be preserved during mitosis (reviewed in (Wang and Higgins 2013)). However, the mitotic histone modification levels were mostly assessed globally (*e.g.,* by methods such as western blots, mass spectrometry, or immuno-staining), and therefore lack spatial genomic information, with few exceptions where ChIP experiments demonstrated a mitotic preservation of histone acetylations at the promoters of several transcribed genes (Kouskouti and Talianidis 2005; Valls et al. 2005). An initial clue regarding the genome-wide organization of histone modifications was achieved by employing ChIP-seq to map H3K4me3, H3K27ac, and H2A.Z in interphase and mitotic cells. Interestingly, the genome-wide maps of H3K4me3 and H2A.Z revealed that the general genomic localization of these marks is conserved upon mitosis entry (Kelly et al. 2010; Liang et al. 2015). Conversely, most H3K27ac genome-wide studies describe a degree of locus specificity in the maintenance of H3K27ac during mitosis (Hsiung et al. 2016; Li et al. 2017; Liu et al. 2017a; Liu et al. 2017b).

Intriguingly, while the genomic localization of H3K4me3 and H2A.Z appears to be highly concordant between mitosis and interphase, the patterns of these histone marks were shown to dramatically differ between mitosis and interphase in specific genomic areas termed NDRs (nucleosome depleted regions) (Kelly et al. 2010; Liang et al. 2015). NDRs are loci highly depleted of nucleosomes, and hence differ from the mostly uniform distribution of nucleosomes across the genome (Lee et al. 2004). NDRs are mainly located within promoters, enhancers, and insulators, and thus are enriched with RNA polymerase and transcription factor binding sites (Fu et al. 2008; Henikoff 2008; Tsompana and Buck 2014). Promoter associated NDRs were shown to disappear during mitosis and contain a nucleosome harboring H3K4me3 (Liang et al. 2015) and H2A.Z (Kelly et al. 2010). However, the histone composition of the nucleosome that occupies these NDRs, as well as the NDR occupation mechanism remain unclear.

To better understand the role of mitotic histone modifications and nucleosome depositioning, we investigated the global dynamics and genomic organization of histone modifications in mitosis and in interphase. Using HeLa-S3 cells as a model system, our study incorporated a synchronization approach that enables the collection of relatively pure preparations of mitotic cells; proteomics analysis focused on quantitative evaluation of the global levels of histone modifications; and finally, systematic genome localization mapping of RNA polymerase II (RNA PolII) and six key histone modifications (H3K4me1, H3K4me3, H3K9ac, H3K27ac, H3K27me3, and H3K36me3).

Our results demonstrate that the histone modification levels and patterns are generally concordant between interphase and mitosis, with one main exception: while histone methylations remain highly similar, histone acetylations show a general reduction in mitosis. Our results expand on previous observations and demonstrate that the epigenomic landscape can serve as a major component of the mitotic bookmarking process. Finally, analysis of NDRs associated with transcription start sites (TSSs), putative enhancers, and putative insulators revealed that the nucleosome that occupies the NDR during mitosis is distinguishable from the surrounding nucleosomes and is characterized by a distinctive set of histone modifications. We propose that the nucleosome adjacent to the interphase NDR (*e.g.*, the +1 nucleosome for TSSs) shifts to the NDR during mitosis and undergoes deacetylation or other modification regulated by HDACs. Small molecule inhibition of HDACs reverts the observed pattern of the shifted nucleosome, lending support to our model. Thus, we provide evidence for a mitotic deposition and modification of the nucleosomes surrounding the NDR of promoters, enhancers, and insulators.

## Results

### Efficient metaphase synchronization of HeLa-S3 cells using a kinesin 5 inhibitor (STC)

Mitotic cells constitute only about 2% of a non-synchronized interphase population. Therefore, determining the chromatin landscape by proteomics and ChIP-seq of this cell cycle phase requires the generation of relatively pure preparations of mitotic cells. The kinesin 5 inhibitor STC (Skoufias et al. 2006) inhibits both separation of the duplicated centrosomes as well as bipolar spindle formation, resulting in cells arrested in metaphase with monoastral spindles (**Figure 1A)**. Following treatment of HeLa-S3 cells with STC we were able to achieve tight synchronization with an average of approximately 90% (SE = 1.6%; n=8) of cells in a metaphase-like stage of mitosis (**Figure 1B**). For simplicity, we refer below to the unsynchronized population as “interphase” cells and to the synchronized one as “mitosis”.

**Figure 1.**
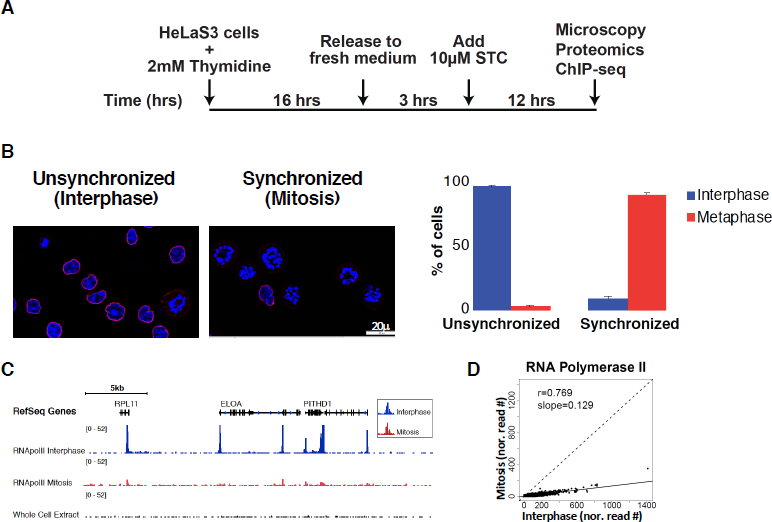
**Efficient metaphase synchronization of HeLa-S3 cells using a kinesin 5 inhibitor (STC) reveals low levels of RNA PolII binding during metaphase. (A)** Schematic representation of the synchronization approach. **(B)** Synchronization was monitored by microscopy using both DAPI (blue) and Lamin B (red) staining. Only cells with the monopolar spindle appearance of the chromosomes and a lack of nuclear envelope were counted as cells in the metaphase stage. The percentage of cells in interphase (blue) and metaphase (red) are shown for both conditions, along with the standard error (SE). For simplicity, we refer to the unsynchronized population as “interphase” and to the synchronized one as “mitosis”. **(C)** Integrative genomic viewer (IGV) (Robinson et al. 2011; Thorvaldsdottir et al. 2013) tracks showing the alignment for RNA PolII ChIP-seq results on an approximately 180Kb region on chromosome 1. The image demonstrates reduced levels of RNA PolII binding in the metaphase sample (red) compared to the interphase sample (blue). **(D)** Scatter plots showing the coverage of RNA PolII peaks in mitosis versus interphase after normalization to 1×10^7^ reads.

### RNA PolII binding is maintained during mitosis at low levels

In accordance with previous studies (Parsons and Spencer 1997), we observed a dramatic reduction in RNA PolII occupancy in the mitotic samples (**Figure 1C-D**). Nevertheless, in both interphase and mitotic samples, RNA PolII binding was almost exclusively at promoters (**Figure S1**). Recent findings suggest that the cell’s transcription pattern is largely retained at a low level throughout mitosis (Palozola et al. 2017). However, we cannot rule out the possibility that the low peaks we observe in mitosis are due to the interphase cells that constitute approximately 10% of our mitotic samples.

### Identification of global changes in histone modifications via mass-spectrometry based proteomic analysis

To gain insight into the global changes in histone modifications during mitosis, we employed a quantitative mass-spectrometry (MS) based approach (Jaffe et al. 2013). By implementing synthetic peptide standards, this MS approach provides high reproducibility and is ideal for comparing histone modification levels between mitosis and interphase. Using this method, we profiled the patterns of histone modifications in three biological replicates of mitotic and interphase samples (**Figures 2** and **S2**). In line with previous observations (Hendzel et al. 1997; Van Hooser et al. 1998; Johansen and Johansen 2006), the H3S10ph modification was highly enriched in the mitotic samples compared to interphase (∼17 fold; p-value < 10^−3^; two sided t-test; **Figure 2**), lending additional support to our synchronization approach. We further estimated the absolute occupancy percentage of individual histone marks by comparing mass spectral intensity vs. known concentrations of internal standards and summarizing the results. While the global levels of histone methylations were mostly similar between mitosis and interphase, histone acetylations as well as K18 ubiquitination were reduced in mitotic samples (∼3-6 fold; p-value < 0.015; two sided t-test; **Figure 2).**

**Figure 2.**
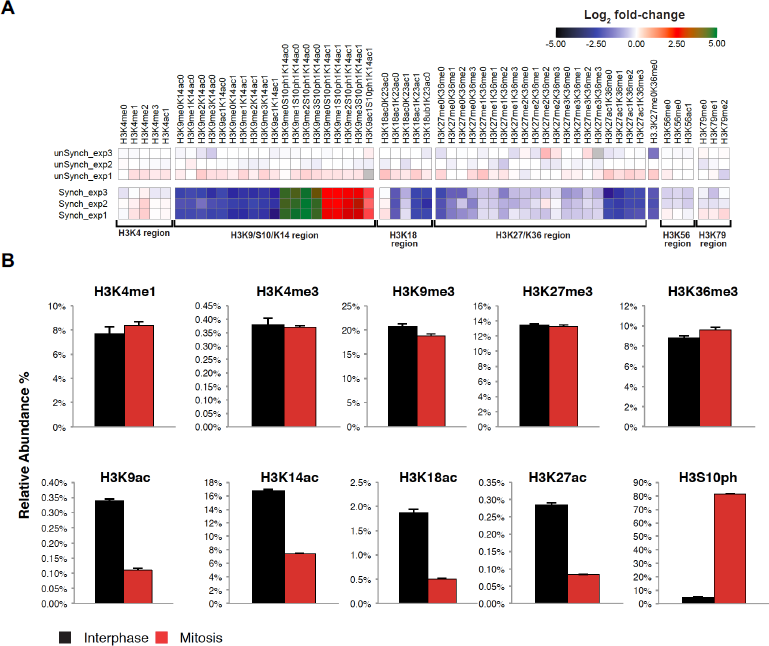
**Identification of global changes in histone H3 modifications by quantitative targeted mass spectrometry analysis. (A)** Heat map (Log2 of fold change) of the different histone modifications as detailed on top. “Regions” (all peptides associated with specific modified/unmodified residue/s) are shown on the bottom. The peptides from these regions were then used to calculate the occupancy percentage of each specific histone modification presented in **B**. Data was normalized to the average signal of interphase samples. **(B)** Relative abundance of selected modifications on H3 tails from interphase or mitotic HeLa-S3 cells. The data was obtained by quantitative targeted mass spectrometry analysis. Each bar represents the percentage of the H3 peptides with the indicated modifications within the total H3 tail peptide population. Note the high levels of the mitosis unique modification, H3S10ph (∼17 fold increase, p-value < 10^−3^; two sided t-test), providing evidence supporting the tight synchronization of the mitotic samples. Similar results were obtained for H4 modifications (**Figure S2**). Note the general decrease in the level of all forms of the K27/K36 peptide in mitosis. This is most probably a result of the mitosis-associated phosphorylation of serine 28 (S28), which is located on the same peptide, reducing the abundance of the non-phosphorylated forms of this peptide in mitosis. We did not include the S28 phosphorylation in our analysis due to lack of an appropriate standard peptide.

We confirmed the distinction between histone methylation and acetylation by performing quantitative immunofluorescence and measuring the mitotic and interphase levels of H3K9ac and H3K4me3. While the levels of H3K4me3 were almost identical in mitosis and interphase as measured by immunofluorescence, H3K9ac levels were reduced in mitosis by 2.7 fold (p-value < 10^−23^; t-test; **Figure S3)**. Immunofluorescence measurements alone are not sufficient for accurate quantification of modification levels since antibody binding during mitosis may be influenced by chromatin condensation or H3S10 phosphorylation. Nevertheless, obtaining similar results from two independent techniques, *i.e.*, immunofluorescence and mass spectrometry, strongly supports the conclusion that there is a key distinction between the protein level dynamics of histone acetylation versus methylation in the transition from interphase to mitosis. However, these results do not show how this difference is reflected spatially in the genomic organization of these histone modifications.

### Analysis of the epigenomic landscape reveals high concordance between mitotic and interphase localization of histone marks

To follow up on the observation from the proteomics analysis and compare the genomic localization of histone marks between mitosis and interphase, we performed ChIP-seq focusing on key histone modifications. We chose six major histone modifications that together capture main chromatin features including active promoters (H3K4me3, H3K9ac, and H3K27ac), putative enhancers (H3K4me1 and H3K27ac which is also associated with active promoters), gene bodies (H3K36me3), and repressed regions (H3K27me3) (Zhou et al. 2011) (**Figure 3**). Surprisingly, we found no major changes between the overall epigenomic landscape of the mitotic and interphase samples, in two biological replicates (**Figures 3A** and **S4**). While this observation was true for all histone modifications examined, there is a clear distinction between methylation marks, which seem to remain intact (with one exception discussed below) and acetylation marks, which show a universal and uniform signal reduction in mitosis (**Figure 3B**). This is in line with our proteomics and immunofluorescence observations regarding the global levels of the modifications in these cell cycle phases.

**Figure 3.**
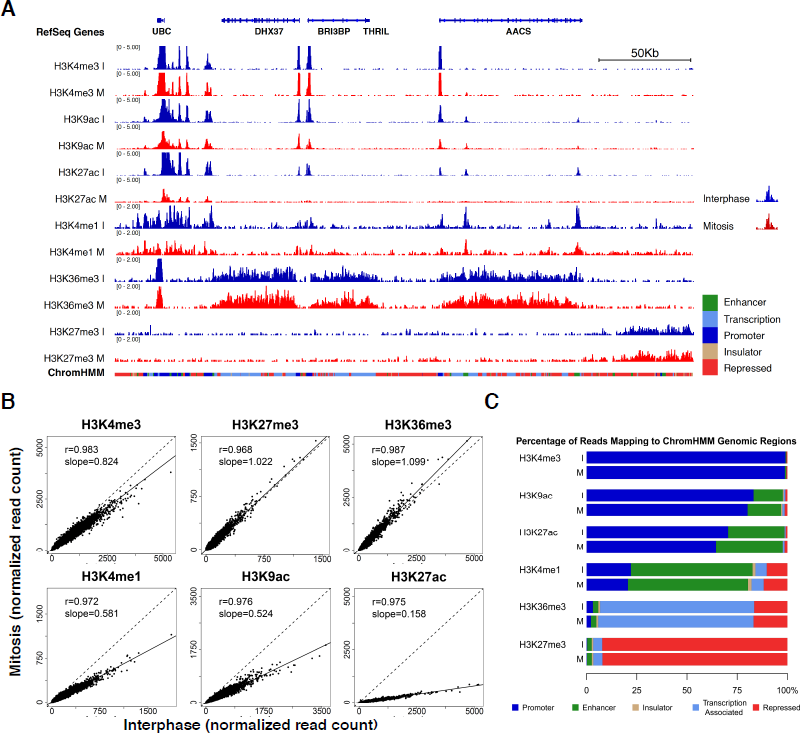
**High concordance between mitotic and interphase histone modification patterns. (A)** Integrative genomic viewer (IGV) (Robinson et al. 2011; Thorvaldsdottir et al. 2013) tracks showing the alignments for the indicated histone modification ChIP-seq results for interphase (blue) and mitosis (red) samples on an approximately 315Kb region on chromosome 12. The scale of each track was adjusted to the total number of reads using the Normalize Coverage Data option in IGV (see methods). The ChromHMM annotation for each genomic region is shown below the plot using the same color code as in **C**. **(B)** Scatter plots showing the normalized read counts for all regions enriched in each modification (see methods for the peak calling details) in mitosis versus interphase. It should be noted that ChIP-seq based quantification may be less accurate than quantification based on mass spectrometry data due to inherent noise in the ChIP-seq method. Thus, in cases of a discrepancy, *e.g.*, H3K4me1, we rely on the mass spectrometry results for quantification, and on the ChIP-seq results for localization. **(C)** Bar plots showing the percentage of reads in peak regions per ChromHMM category.

The only exception to our above observation is the ChIP-seq data for H3K4me1. The signal for this histone modification is moderately reduced in mitosis (**Figure 3B**). However, since the proteomics analysis displays a high concordance between the levels of H3K4me1 in interphase and mitosis (even showing a slight increase in mitosis), the discrepancy might be related to the challenges in quantifying ChIP-seq signal due to the inherent noise in this method. We thus relied in this case on the MS data for assessing the levels of H3K4me1, and on the ChIP-seq data regarding the localization of this histone modification.

To quantify the genomic changes in the levels of histone modifications between mitosis and interphase, we used the ENCODE mappings of chromatin environments derived from the combined ChromHMM annotations (Ernst and Kellis 2012). We compared the organization of histone modifications within the chromatin environments and found that the distribution of the methylation signals in interphase and mitosis was almost identical. Conversely, the genomic distribution of the H3K27 and H3K9 acetylations differed slightly between these cell cycle phases. Both acetylations presented an increase in signal over enhancer regions (5% and 2%; respectively), and concurrent reduction in promoter regions (6% and 3%; respectively) in mitosis (**Figure 3C**).

### The nucleosome depleted regions (NDRs) become occupied during mitosis by a nucleosome distinguishable from its neighbors

Previous reports demonstrated that nucleosome depleted regions (NDRs) associated with the transcription start site (TSS) of promoters are lost during mitosis, and the gap is occupied by a nucleosome containing H3K4me3 and H2A.Z (Kelly et al. 2010; Liang et al. 2015). To better understand this mitotic change in the chromatin structure, we leveraged the variety of histone modifications we mapped in mitosis and interphase to study the histone modifications of the nucleosome that occupies the NDR during mitosis. As nucleosome depleted regions were also observed within enhancers and insulators (Henikoff 2008; Tsompana and Buck 2014), the range of marks we profiled enabled the deep investigation of NDRs associated with promoters, putative enhancers, and insulators for the first time.

First, we focused on H3K4me3, which was previously shown to occupy promoter NDRs during mitosis (Liang et al. 2015). In line with Liang *et al.*’s report, the datasets we generated were mostly devoid of the promoter NDR structure in mitosis (**Figure 4A**), even though our samples were synchronized using a different approach (STC vs. nocodazole). Additionally, extension of this analysis to include putative enhancers and insulators revealed that in mitosis, a nucleosome containing methylated histones also occupies NDRs associated with these regulatory regions (**Figures 4A** and **S5 – S7**).

**Figure 4.**
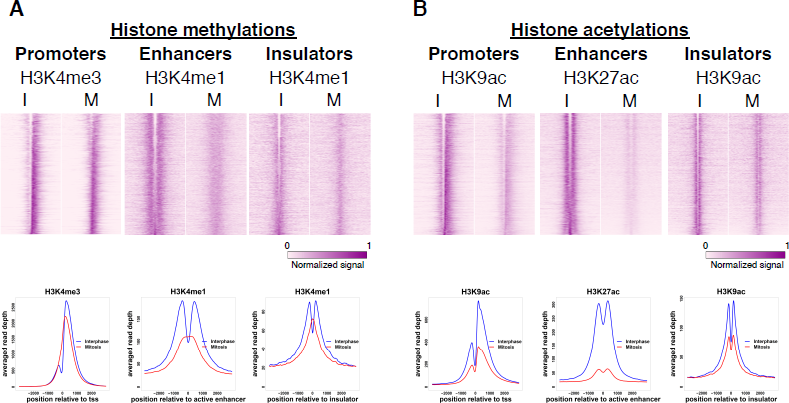
**The nucleosome that enters the NDR during mitosis is distinguishable from the surrounding nucleosomes.** Top: heat maps showing the normalized and standardized read depth at 6Kb regions centered at either TSSs, enhancers, or insulators, for H3K4 methylations (n=6488, 7062, 4367; respectively. (**A)**), and H3K9 or H3K27 acetylations (n=6808, 5565, 3033; respectively. (**B**)). Bottom: “Metagene plots” showing the average read depth for all regions in each category are under the relevant heat map. Similar plots for additional modifications and for a biological repeat are shown in **Figures S5-S7**.

Surprisingly, when we analyzed the organization of the histone acetylations H3K9ac and H3K27ac over NDRs, the binding structure was mostly preserved during mitosis as opposed to the pattern observed for the histone methylations, suggesting that the nucleosome that occupies the NDR is deacetylated (**Figures 4B** and **S5-S7**). To corroborate our observation, we analyzed data from G1E murine erythroblast cells (Hsiung et al. 2016). Consistently, in G1E murine cells the nucleosome that occupies the NDR is mostly deacetylated (**Figure S8**), indicating that this phenomenon is conserved between species and cell types. Altogether, these data demonstrate that the nucleosome that occupies NDRs during mitosis is methylated but not acetylated, hence it is distinguishable from the neighboring nucleosomes which are dually marked by methylation and acetylation.

### Classification of the NDRs according to nucleosome modifications during mitosis

To further characterize the nucleosome that occupies TSS associated NDRs during mitosis, we looked at the coverage profiles of genes marked with H3K4me3, H3K9ac, and H3K27ac at their TSSs, and sorted the profiles by occupancy of mitotic H3K9ac at the TSS. The sorted profiles revealed four gene groups based on the combination of modifications occupying the NDR (**Figures 5A-B** and **S9**). The groups were classified as follows: *Group 1*: retention of the NDR during mitosis. *Group 2*: loss of the NDR during mitosis due to the entry of a nucleosome with solely methylated histones. *Group 3*: loss of the NDR during mitosis due to the entry of a nucleosome with both methylated and acetylated histones. *Group 4:* absence of the NDR in both interphase and mitosis. We disregarded *Group 4* in further analyses since it already lacked the NDR structure in interphase. Of the promoters analyzed, we found that approximately 22% belong to *Group 1*, 61% to *Group 2*, and 9% to *Group 3*.

To further assess these promoters, we first characterized the associated genes. We found that of the genes in each of *Groups 1-3,* 88-92% were shown to be transcribed in HeLa cells (Hart et al. 2013), 5% are cell cycle genes (as defined by (Whitfield et al. 2002)), 59-61% are housekeeping genes, and 13-15% are tissue specific genes (as defined by (Uhlen et al. 2015)) (**Table S1**). Second, we compared the percentage of GC in each group and found that *Group 1* has a higher GC content than *Groups 2* and *3* (approximately 70% vs 62% and 60% in *Groups 2* and *3*, respectively; **Figure 5C**). Thus, the GC content appears to distinguish between regions that retain the NDR during mitosis and regions that lose it, regardless of the histone modifications on the NDR-occupying nucleosome. As GC content may be associated with the binding of specific transcription factors (TFs), we next employed DREME, a discriminative motif discovery tool (Bailey 2011), to identify motifs enriched in *Group 1* relative to the other groups. Indeed, along with the high GC content observed in *Group 1*, many of the motifs were enriched for GC (**Figure S10).**

To further investigate the characteristics of these three groups, we studied the binding patterns of TFs mapped by ENCODE in HeLa-S3 cells and our RNA PolII occupancy data from interphase cells. Similar to the GC content distinction, RNA PolII as well as most of the TFs (*e.g.*, HCFC1, BRCA1, and MXI1; **Figures 5D** and **S11**) demonstrated a binding pattern that differed between *Group 1* and the two other groups (*2* and *3*), whereas only a limited number of factors (*e.g.,* E2F1 and USF2; **Figures 5D** and **S11**) show distinct binding patterns for each group. This observation, together with the GC content results, suggests that there is an intrinsic division between genes that have a nucleosome in the NDR during mitosis versus genes that do not.

Next, to better understand what distinguishes *Group 2* from *Group 3*, we evaluated the association of HDAC binding to each of these groups. Analysis of our and public HDAC2 and HDAC6 ChIP-seq data from unsynchronized cells (Wang et al. 2009; Ram et al. 2011) revealed a statistically significant difference between these groups: 43% of the TSSs in *Group 2*, whose NDR-occupying nucleosome is deacetylated, overlapped with an HDAC peak, while only 35% of the TSSs in *Group 3*, whose NDR-occupying nucleosome is acetylated, were occupied by an HDAC (p-value ∼ 0.001; Chi squared test). Thus, it appears that the presence of HDAC may contribute to the observed difference in the acetylation patterns.

Finally, to identify a potential function for the differential NDR patterns, we analyzed gene reactivation kinetics following mitotic exit using a recently published dataset (Palozola et al. 2017). We found that genes in *Groups 1* and *2* are reactivated significantly earlier than the genes in *Group 3* (**Figure 5E** p-value = 0.00235; ANOVA). This finding suggests that the NDR nucleosome association pattern may differentiate between genes that are important promptly following mitotic exit versus genes that are only necessary later. Indeed, gene ontology analysis of the genes associated with the three Groups revealed that *Groups 1* and *2* are enriched for genes which may be relevant immediately upon mitotic exit such as genes involved in cell signaling, localization, and RNA processing (**Figure S13**). These results suggest that the NDR pattern seen at the promoters of genes in *Groups 1* and *2* may be important for the rapid reactivation of transcription following mitosis.

### Mitosis associated nucleosome deposition into the NDR

Nucleosome occupancy at NDRs during mitosis was previously explained as a consequence of the penetration of the +1 nucleosome into the NDR (Kelly et al. 2010). However, the absence of acetylation of this nucleosome challenges this model. We envisioned two possible models for the integration of the nucleosome into the NDR during mitosis. The *first model* being the incorporation into the NDR of a new nucleosome whose histones are methylated but not acetylated. The *second model* proposes, as postulated previously, that the surrounding nucleosomes shift into the NDR in mitosis, and the histones are actively deacetylated by HDACs (histone deacetylases).

To discriminate between the two models, we first evaluated the dynamics of H3K4me3 in interphase and mitotic samples. H3K4me3 was shown both by a previous study and our analysis to occupy the NDR during mitosis. To evaluate nucleosome movement from our data, we measured the difference in the H3K4me3 signal in mitosis compared to interphase for promoter NDRs (Delta plots; see **Methods**). We observed that the increase in nucleosome occupancy at the TSS is followed by a decrease in nucleosome occupancy in the downstream region, suggesting a shift of the +1 nucleosome into the NDR. This observation is in line with Kelly *et al.*’s findings (Kelly et al. 2010). Furthermore, we identified similar patterns of surrounding nucleosomes entering NDRs during mitosis at putative enhancers and insulators (**Figures 6A** and **S14**).

Next, to further provide support for the second model, we treated interphase and mitotic cells with the global HDAC inhibitor TSA. Perturbation of HDAC activity eliminates the general mitotic hypoacetylation, and ChIP-seq performed on perturbed samples demonstrates highly similar H3K9ac signal intensities in interphase and mitosis (**Figure 6B** and **S14**). Additionally, inhibition of HDAC activity altered the histone acetylation pattern we observed previously at the NDRs, and in TSA-treated mitotic cells the NDR-occupying nucleosome has acetylated histones (**Figure 6C** and **S14**). Delta plots demonstrated that the increase in the level of acetylated histones at the NDR is accompanied by a decrease in the signal at the surrounding regions, which was indiscernible in untreated cells due to global mitotic deacetylation (**Figure 6D** and **S14**). This further supports the notion that this nucleosome shifts into the NDR from a surrounding position. A similar observation was seen in enhancer and insulator regions (**Figure S15**). Finally, H3K4me3 patterns were not affected by TSA treatment, thus minimizing the possibility of a global effect (**Figure S16**).

## Discussion

In this study, we aimed to gain a better understanding of the mitotic bookmarking process and the organization of chromatin during this phase. We designed an interdisciplinary approach that merges proteomics, immunofluorescence, automated ChIP-seq, and computational analysis that allowed the robust identification of changes between the interphase and mitotic chromatin organization. We incorporated the results of biological replicates for all measurements to obtain quantitative information regarding the global amounts of modified histones in each cell cycle phase. This was achieved by employing MS and immunofluorescence, and by generating genome-wide chromatin maps for RNA PolII and six key histone modifications. These epigenomic landscapes provide insight into the spatial chromatin organization during interphase and mitosis. To the best of our knowledge, we are the first to conduct a systematic molecular survey that combines global (proteomics and immunofluorescence) and spatial (epigenomics) measurements in interphase and mitosis.

Mass-spectrometry (MS) data yields reliable quantitative results and therefore can be used to measure global changes in histone modification levels. Additionally, ChIP-seq data, while providing lower-accuracy quantitative measurements, contributes valuable spatial information. The incorporation of MS data is particularly valuable when assessing histone acetylation levels, since mitosis-specific H3S10 and H3S28 phosphorylations may limit the functionality of H3K9ac and H3K27ac antibodies (**Figure S17**) (Rothbart et al. 2015; Cornett et al. 2017). For this reason, we rely on the MS results to evaluate the mitotic levels of these acetylations, rather than on the ChIP-seq data. Nevertheless, the spatial information provided by the ChIP-seq results is reliable for all modifications, assuming a genome-wide uniform distribution of the mitotic phosphorylations. The integration of these varying approaches has allowed us to gain insight into both the quantification and localization of each tested histone modification.

All of our results (excluding the immunofluorescence experiments) are based on a prolonged mitotic arrest. Although this approach is widely used in the field (Kadauke et al. 2012; Hsiung et al. 2015; Hsiung et al. 2016; Palozola et al. 2017), we cannot rule out the possibility that some of our observations may be a consequence of this abnormal mitosis phase. It should be noted that despite the prolonged mitotic arrest, the majority of cells retained the ability to successfully reenter the cell cycle (**Figure S18).** Further method development is needed in order to perform similar experiments on unperturbed mitotic cells.

Analyzing the proteomic and epigenomic datasets provided several major insights. We observed that although mitotic chromatin is highly condensed, global levels of histone methylations are mostly maintained. In contrast, histone acetylation abundance was reduced in mitotic samples as seen by measuring both global protein levels (via proteomics and immunofluorescence) and DNA association (via ChIP-seq). Yet, the change in acetylation levels between mitosis and interphase was moderate (about 2-6 fold) compared, for instance, to the variation observed in H3S10ph levels (about 17 fold) between these two phases (**Figure 2**). Furthermore, comparison of the genomic localization between interphase and mitotic samples demonstrated that the organization of these modifications was tightly maintained. Altogether, apart from the patterns we observed in NDRs and discuss below, we did not identify local differences in the organization of the histone modification patterns between mitosis and interphase, suggesting that the genome undergoes global, uniform changes during mitosis.

The observation that histone modification patterns are concordant between interphase and mitosis lends support to the hypothesis that the preserved genomic organization of these marks plays a bookmarking role. This model is an important addition to our understanding of the epigenomic memory during the cell cycle. For many years mitotic chromosomes were considered to be transcriptionally silenced (Taylor 1960; Prescott and Bender 1962; Parsons and Spencer 1997), devoid of most transcription factors (Martinez-Balbas et al. 1995; Gottesfeld and Forbes 1997; Prasanth et al. 2003), and condensed (Antonin and Neumann 2016). Thus, it was unclear how epigenomic information is transmitted throughout the cell cycle. Initial studies have demonstrated that bookmarking is carried out by a limited number of transcription factors that are retained on the chromosomes during mitosis to enable the reestablishment of the transcriptional program in G1 (Kadauke et al. 2012; Kadauke and Blobel 2013).

In contrast, our study suggests that the overall epigenomic structure in interphase is vastly maintained during mitosis. This observation is in line with several recent studies, which indicate various levels of similarity between the mitotic and interphase chromatin structure. First, it was shown both in drosophila and mice that a large portion of the transcription machinery, including many transcription factors and chromatin regulators, remain on the chromosomes during mitosis (Black et al. 2016; Teves et al. 2016; Liu et al. 2017b). Second, a genome-wide DNase-seq experiment revealed that genome accessibility is widely preserved and only locally modulated during mitosis (Hsiung et al. 2015). Finally, a recent study utilized 5EU labeling to demonstrate that many genes exhibit transcription during mitosis (Palozola et al. 2017). Taken together, these, as well as our observations suggest that despite their condensation, mitotic chromosomes retain a base level of their original chromatin structure and are partially involved in active transcription. This can explain the reestablishment of transcription after mitosis without assuming additional bookmarks.

Recently, several studies employed ChIP-seq to assess histone modification levels during mitosis. There are however, conflicting observations regarding the direction and uniformity of the changes in mitotic H3K27ac. While we observed a uniform decrease in H3K27ac levels in mitosis relative to interphase (**Figures 2** and **3B**), a recent study in breast cancer MCF-7 cells saw a uniform increase in the levels of this modification in mitotic cells (Liu et al. 2017a). Other studies have demonstrated that the change in H3K27ac levels in mitosis is heterogeneous (Hsiung et al. 2016; Liu et al. 2017b). We cannot rule out the possibility that these variations are a consequence of technical differences (*e.g.*, synchronization techniques) between the studies. However, the variations may also reflect a biological distinction between cancer or fully differentiated cells, and cells that carry the potential for further differentiation. In line with this notion, mitosis constitutes a window of opportunity for altering cellular fates via mitotic chromatin reprogramming (Halley-Stott et al. 2014). Thus, H3K27ac, a modification linked to cell fate decisions (Long et al. 2016), may exhibit a heterogeneous change in profile in cells possessing the potency to differentiate (ESC and G1E erythroblasts), and a uniformly changing profile in cancer cells (HeLa-S3 and MCF-7).

While we observed retention of the epigenomic organization during mitosis both for histone methylations and acetylations, there was a uniform genome-wide deacetylation of histone H3K9 and H3K27. Global deacetylation of histones during mitosis has been previously observed, but its precise role in mitosis is not fully understood. An early study revealed that blocking deacetylation by HDAC inhibition shortly before mitosis in human primary fibroblasts causes severe mitotic defects (Cimini et al. 2003). It has also been shown that independent of its role in transcription regulation, histone deacetylation during mitosis plays a structural role in mediating Aurora B-dependent phosphorylation of mitotic chromosomes, which is important for their condensation (Li et al. 2006). On the other hand, other studies have suggested a transcription dependent mechanism (Noh et al. 2009), which would presumably be locus specific. Thus, our results, which show a uniform genome-wide deacetylation of H3K9 and H3K27, strongly support the structural role of the deacetylation over a transcription dependent mechanism. Indeed, blocking histone deacetylation by TSA strongly reduces the cells’ ability to exit mitosis following STC release (**Figure S18**).

Previous studies demonstrated that during mitosis there is a shift of the +1 nucleosome into the NDR that is associated with transcription start sites (Kelly et al. 2010; Liang et al. 2015). Yet, as these studies focused either on H2A.Z (Kelly et al. 2010) or H3K4me3 (Liang et al. 2015), there was no data regarding other components of the nucleosome that occupies the NDR in mitosis. Furthermore, it was unclear whether this phenomenon is restricted to NDRs associated with promoters, or if it is global and can also occur in regions associated with enhancers or insulators.

Leveraging the comprehensive paired datasets we generated for interphase and mitosis, we were able to expand upon these initial observations and study a broad set of histone modifications, as well as the NDRs associated with enhancers and insulators. We observed that while our H3K4me3 data exhibited very similar patterns to the ones seen by the Shilatifard group (Liang et al. 2015), the nucleosome occupying the NDR in mitosis was devoid of histone acetylations H3K9ac and H3K27ac. This is a very intriguing observation, as it implies that the nucleosome that enters the NDR during mitosis differs from its neighboring nucleosomes, which comprise H3K4me3, H3K9ac, and H3K27ac. By analyzing public H3K27ac data from murine erythroblast cells, we showed that this NDR structure is also found in mouse cells, suggesting that the acetylation pattern we observed is evolutionarily conserved. Alternatively, the reduction in the acetylations of the nucleosome that enters the NDR during mitosis that we observed may be a result of the mitosis-specific phosphorylation of nearby residues (Raynaud et al. 2014), which can hinder the ability to detect acetylations (**Figure S17**). Thus, the observed pattern may in fact reflect a specific phosphorylation of the moving nucleosome, rather than a deacetylation. Further research is needed in order to understand the cross-talk between acetylation and phosphorylation during mitosis.

The role of the mitotic movement of the nucleosome into the NDR is not clear. It may be a mere consequence of the eviction of RNA polymerase and other factors that frees up the DNA at the NDR to become occupied by adjacent nucleosomes. However, the specific deacetylation that takes place precisely on the histones that move into the NDR likely requires a designated mechanism, evidence that this process is biologically important. Indeed, the transcription reactivation kinetics differ between genes in *Groups 1* and *2* versus *Group 3* (**Figure 5E**). This suggests that the type of nucleosome that populates the NDR, deacetylated (*Groups 1* and *2*) or acetylated (*Group 3*), may affect the reactivation kinetics.

An appealing explanation for this finding is that the deacetylation serves to bookmark the region as an NDR in interphase and enables the removal of the penetrating nucleosome immediately following mitosis. In line with this hypothesis, it has been shown that some chromatin remodeling complexes are affected by histone modifications (Swygert and Peterson 2014). Thus, the deacetylation may facilitate the removal of the nucleosome entering the NDR by recruiting specific remodeling complexes after the mitosis phase. Further studies are needed to evaluate this hypothesis and investigate whether nucleosome repositioning is indeed important for the bookmarking process.

Following our observation that the nucleosome that penetrates the NDR during mitosis has a distinctive composition of modifications, we have shown that the deacetylation of this nucleosome is HDAC dependent (**Figure 6**). However, the exact mechanism that localizes the deacetylation specifically to this nucleosome remains unclear. We and others have previously shown that HDACs bind tightly at the TSS (Wang et al. 2009; Ram et al. 2011). This suggests that HDAC positioning may play a role in the specific deacetylation of the shifting nucleosome at this region. Alternatively, the shifting nucleosome may be more dynamic than the others, thus during its partial disassociation from the DNA it can become deacetylated. Indeed, it has been shown that histones associated with promoter and regulatory regions are less stable than histones localized at other genomic loci (Jin and Felsenfeld 2007; Jin et al. 2009).

Not all NDRs undergo the same mitosis-associated changes. We found that in most cases the penetrating nucleosome is indeed deacetylated, however there is a large group of promoters in which no nucleosome penetrates the NDR during mitosis (*Group 1*; **Figure 5**). Interestingly, compared to the rest of the promoters, *Group 1*’s promoters are distinguishable already during interphase in several ways: (***i***) these regions have a higher GC content; (***ii***) the binding of many TFs is less localized to the TSS of these promoters (**Figure 5**); and (***iii***) the NDR seems to be wider (see for example the H3K4me3 plot in **Figure S12**).

Taken together, these findings suggest that basic differences in the promoter architecture reflect the mechanism that preserves the NDR during interphase. Only some of the mechanisms associated with these different promoter architectures are active during mitosis, and this may explain the differential loss of some NDRs. Thus, for instance, sequence based mechanisms should be preserved during mitosis, whereas interphase NDRs that are kept via the binding of transcription factors (Struhl and Segal 2013), may be lost as cells undergo mitosis. Further work is needed to understand these mechanisms and their importance in mitotic function.

Overall, together with several recent studies, our work provides an important cornerstone necessary for shifting the paradigm that has lingered for many years. We can reason that while structurally the chromatin goes through major changes during mitosis, molecular characteristics such as the organization of histone modifications are maintained, even if their levels may be reduced. This provides a simpler explanation for the restoration of chromatin structure following mitosis.

## Methods

### Cell Culture and synchronization

HeLa-S3 cells were grown in Spinner flasks in Dulbecco’s modified Eagle’s medium (DMEM) supplemented with 10% Fetal Bovine Serum (FBS), Penicillin-Streptomycin, L-Glutamine, Sodium Pyruvate and 0.1% Pluronic F-68. Cells were incubated at 37^°^C and 5% CO_2_. Cells were pre-synchronized in G1/S by addition of 2mM thymidine for 16 hours, washed with PBS, released for 3 hours in fresh medium and arrested with S-Trityl-L-cysteine (STC) (164739, Sigma) for 12 hours. HDAC blocking was done with 150nM Trichostatin A (TSA, T1952, SIGMA) which was added either to the unsynchronized culture or 3 hours after adding the STC, for 9 hours.

### Immunofluorescence and Microscopy

Cells were fixed with either 70% ethanol or 4% paraformaldehyde, blocked with 5% BSA, and stained either with mouse anti LaminA (1:1000; L1293, Sigma) followed by donkey anti-mouse DyLight^®^ 550 (1:400; ab98795, Abcam), or with rabbit anti H3K9ac (1:1000; C5B11, Cell signaling) or H3K4me3 (1:1000; C42D8, Cell Signaling Technologies) followed by Alexa fluor 488 goat anti rabbit IgG(H+L) (1:400; A11011, Invitrogen). Cells were counterstained with DAPI.

### Proteomics

Three replicates of HeLa-S3 cells (either unsynchronized or STC synchronized) were harvested. Exactly 2.1 million cells from each sample were snap-frozen in liquid nitrogen and stored at −80^°^C. Global chromatin profiling (GCP) of cell pellets to quantify the levels of histone modification was performed as described previously (Creech et al. 2015). In brief, cells were lysed, nuclei were isolated, histones were extracted by sulfuric acid and were then precipitated by trichloroacetic acid. Next, the samples were propionylated, desalted, and digested with trypsin overnight. Samples were subjected to a second propionylation, and then desalted using a C18 Sep-Pak Cartridge (Waters). Prior to MS analysis, a mix of isotopically labeled synthetic peptides was added to each sample. Next, the peptides were separated on a C18 column (15 cm x 75 um ID, ReproSil-Pur 120 C18-AQ, 1.9 µm) using an EASY-nLC 1000 (Thermo Fisher Scientific) and analyzed using the GCP assay (a parallel reaction monitoring targeted LCMS method) using a Q Exactive™-plus mass spectrometer (Thermo Fisher Scientific) as described previously (Creech et al. 2015).

The GCP employs high mass accuracy (∼3 parts per million) / high resolution (R= 17,500 at *m/z* 400) mass spectrometry, thus it enables distinguishing between an adduct of trimethylation (42.0470 Da) from acetylation (42.0106 Da).

Standard operating procedures (SOPs) for the GCP assay, including synthetic peptide master mixture formulation, can be found at: https://panoramaweb.org/labkey/wiki/LINCS/Overview%20Information/page.view?name=sops.

### Chromatin immunoprecipitation followed by sequencing (ChIP-seq)

ChIP-seq was performed either manually on individual samples or automatically using the Bravo liquid handling platform (Agilent model 16050-102, “Bravo”) as described previously (Busby et al. 2016). In accordance with our efforts to promote reproducibility, all antibodies used were monoclonal. The following antibodies were used: anti RNA polymerase II 8WG16 (ab817, Abcam), anti H3K4me1 (D1A9, Cell Signaling Technologies), anti H3K4me3 (C42D8, Cell Signaling Technologies), anti H3K9ac (C5B11, Cell Signaling Technologies), anti H3K27ac (D5E4, Cell Signaling Technologies), anti H3K27me3 (C36B11, Cell Signaling Technologies), anti H3K36me3 (D5A7, Cell Signaling Technologies), and anti CTCF G.758.4 (MA5-11187, Invitrogen). The specificity of all antibodies targeting histone modification were assessed by us (Busby et al. 2016) and others (Rothbart et al. 2015), and the array datasets are publicly available at www.histoneantibodies.com.

### Bioinformatic analysis

ChIP-seq reads were aligned to the human genome (hg19) using either BWA (Li and Durbin 2009) or Bowtie (Langmead and Salzberg 2012). Duplicate alignments were removed with Picard Tools MarkDuplicates (http://broadinstitute.github.io/picard/). Only QC-passed reads were used for the analyses (see **Table S3** for read counts of all sequenced samples). Peak detection on merged replicates (with a whole cell extract dataset as a control) was done using HOMER (Heinz et al. 2010) with default histone style parameters for all histone modifications, and default factor style parameters for RNA PolII and CTCF.

Scatter plots were made from read counts (using bedtools multicov (Quinlan and Hall 2010)) at merged interphase and mitosis peaks and normalized to 10 million total reads. ChromHMM (Chromatin State Discovery and Characterization) genomic annotations were downloaded from the UCSC genome-browser for HeLa-S3 cells (Ernst and Kellis 2013) and combined into five main categories. The ChromHMM distribution plot (**Figure 3C**) shows the percentage of reads in peak regions per ChromHMM annotation.

IGV snapshots show TDF files created from the aligned sequence data (using the count command from IGV tools (Robinson et al. 2011; Thorvaldsdottir et al. 2013)). The scale of each track was adjusted to the total number of reads using the Normalize Coverage Data option in IGV which multiplies each value by [1,000,000 / (totalReadCount)] (**Figures 1C** and **3A**).

For generating coverage profiles, metagene plots, and delta plots, we first fixed the length of each read to the average of the mitosis and interphase fragment length modes. We analyzed the following NDRs – TSSs (using gene coordinates from UCSC (Karolchik et al. 2004)); Enhancers (using regions with H3K4me1, H3K27ac, and UCSC P300 HeLa-S3 peaks (wgEncodeEH001820)); and insulators (using merged interphase and mitosis CTCF ChIP-seq peaks), each containing a unique set of loci. In each set, only regions located > 6Kb from each other, and overlapping modification peaks within 500bp on either side of the NDR, were included in the analysis.

Coverage profiles (heat maps) were created from the read depth (calculated using samtools bedcov (Li et al. 2009)) of each modification in a window of 6kb around the NDR in 25bp bins. Read depth was normalized to 10 million total reads, and the depths of each row were then standardized to the range [0, 1]. Heat maps are sorted by NDR occupancy in interphase.

Combined coverage profiles (**Figures 5A and S9**) only include regions with peaks for all of the modifications of interest overlapping the NDR. The normalized read depth was converted to a z-score for mitosis and interphase of each modification separately, and then each row was standardized (to [0, 1]). Heat maps are sorted by TSS occupancy of H3K9ac in mitosis. Visual inspection of the sorted profiles revealed gene groups based on the combination of modifications at the TSS. Metagene plots were produced by averaging the coverage profiles. Delta plots were made by averaging the result of subtracting the interphase coverage profiles from the mitosis coverage profiles.

For the boxplots (**Figure 5C**), GC content was measured around the TSS (+/-500bp) of genes using bedtools nuc, and corresponding p-values (t-test) and effect sizes were calculated.

HDAC2 and HDAC6 peaks for CD4 (Wang et al. 2009) and K562 cells (Ram et al. 2011) were downloaded and merged. The percentage of genes in each group that overlapped an HDAC peak at the TSS (one bp) was determined and p-values were calculated using a chi-squared test.

ChIP-seq datasets were downloaded from ENCODE (see **Table S2** for ENCODE file names) and metagene profiles of gene groups were created using deepTools plotProfile (Ramirez et al. 2016). Counts at the TSS (20bp on either side) were totaled and corresponding p-values (t-test) were calculated.

Lists of HeLa-S3 transcribed genes (only genes with a zFPKM > −3) (Hart et al. 2013), cell cycle genes (Whitfield et al. 2002), and housekeeping/tissue specific genes (www.proteinatlas.org) were downloaded and the percentage of genes in each group that overlapped each of these datasets was determined.

For the transcription reactivation kinetics analysis (**Figure 5E**), data was downloaded (Palozola et al. 2017) and profiles of the mean for each gene group (after subtracting the value at the first time point from each of the subsequent time points) were plotted. Error bars and shading depict the standard error at each time point. P-value was calculated using a Repeated Measures ANOVA performed on *Groups 1* and *2* combined versus *Group 3*.

GO analysis was done with the GOrilla (Gene Ontology enRIchment anaLysis and visualization) online tool (Eden et al. 2009).

**Figure 5.**
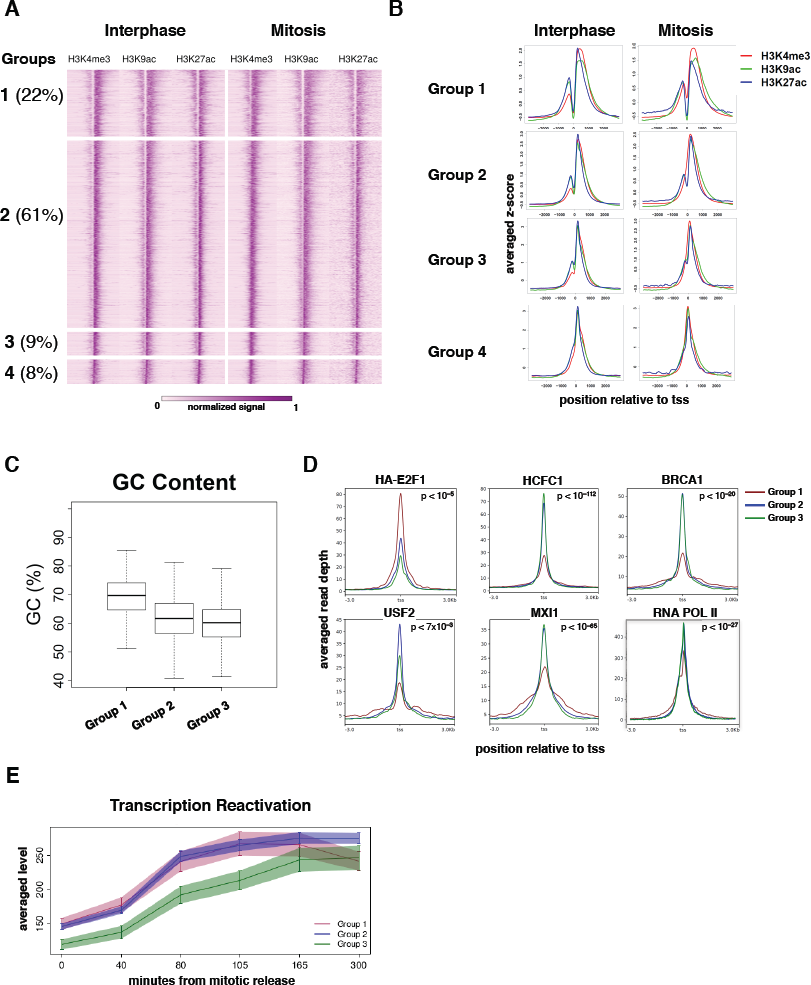
**Characterization of the TSS associated NDRs. (A)** A heat map showing the normalized and standardized z-scores at 6Kb regions centered at the TSS of 5731 genes that are occupied by H3K4me3, H3K9ac, and H3K27ac. The regions were manually divided into four groups (*Groups 1-4*) after visual inspection; the percentage of regions in each group is shown in parentheses. **(B)** Metagene plots showing the average occupancy of each modification for the four groups. **(C)** Boxplot showing the difference in the GC content at the TSS for *Group 1-3*. The difference between *Group 1* and either *Group 2* or *Group 3* is highly significant (p-value = 1.67e-133 and 3.97e-84; t-test, and effect size = 0.88 for *Group 2* and 1.12 for *Group 3*). **(D)** Metagene plots showing ENCODE ChIP-seq data and our RNA PolII data for *Groups 1-3*, p-values were obtained either by comparing *Group 1* to the combination of *Groups 2* and *3* (for HCFC1, BRCA1, MXI1, and RNA PolII), or by determining the maximal p-value from all pairwise comparisons (for HA-E2F1 and USF1). For a comprehensive list of all p-values see **Table S2**. The same analysis for all available ENCODE HeLa-S3 data is shown in **Figures S11-S12**. **(E)** Profiles showing transcription reactivation kinetics following release from mitosis. The difference between *Groups 1* and *2* versus *Group 3* is significant (p-value = 0.00235; ANOVA – note: this tests the change over time of the transcription levels and is therefore not sensitive to the difference in initial level (time = 0) between the groups).

**Figure 6.**
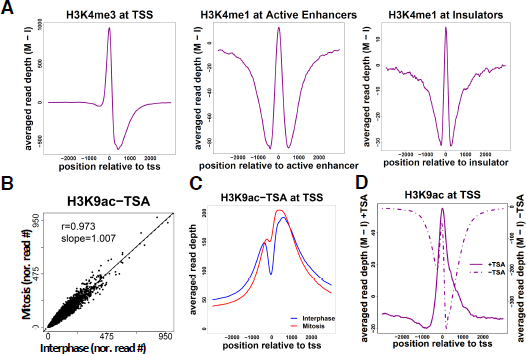
**Nucleosome shifting and deacetylation may explain NDR disappearance. (A)** Delta metagene plots (see methods for details) showing the average read depth of the indicated modifications after subtracting the interphase depths from the mitotic depths. **(B)** A Scatter plot showing the normalized read counts at H3K9ac peaks on cells treated with TSA in mitosis versus interphase. **(C)** Metagene plot for H3K9ac data after TSA treatment. **(D)** Delta metagene plot for H3K9ac data before (**dotted line**) and after (**solid line**) TSA treatment. Similar results were obtained in a biological replicate (**Figure S13**).

## Acknowledgments

We thank Dr. Michele Busby and Dr. Melissa Gymrek for fruitful discussions, Yutong Qiu, Miguel Alcantar, and Michal Schlesinger for technical assistance, and Jason Ernst for assistance with HeLa-S3 ChromHMM data. This research was supported by the Israel Science Foundation (grant no. 184/16 to IS), the Israel Science Foundation – Broad Institute Joint Program (grant no. 1972/15 to IS and AG), the United States-Israel Binational Science Foundation (BSF), Jerusalem, Israel (grant no. 2015272 to IS and AG), and NIH grants (U54 HG008097 to JDJ and MIRA R35GM124736 to SBR).

## Data accessibility

Data can be accessed at the NCBI Gene Expression Omnibus (GEO; http://www.ncbi.nlm.nih.gov/geo/) with accession number GSE108173.

---

**Figure S1. RNA PolII binds almost exclusively at promoters.** Bar plots showing the percentage of reads in peak regions per ChromHMM category for interphase (I) and mitosis (M).

**Figure S2. Identification of global changes in histone H4 modifications by quantitative targeted mass spectrometry analysis. (A)** Heat map (log2 of fold change) of the different histone modifications as detailed on top. Data was normalized to average signal of interphase samples. **(B)** Relative abundance of selected modifications on H4 tails from interphase or mitotic HeLa-S3 cells. The data was obtained by quantitative targeted mass spectrometry analysis. Each bar represents the percentage of the H4 peptides with the indicated modifications within the total H4 tail peptide population.

**Figure S3. Immunofluorescence analysis of H3K9ac and H3K4me3 levels in mitotic samples. (A)** A representative immunofluorescence picture of H3K4me3 and H3K9ac (green) and DAPI (blue) staining. The cell cycle stage was determined from the DAPI image. **(B)** Quantification of the green signal intensity in multiple (N>28) mitotic and interphase cells. Average and standard error are shown. The difference between interphase and mitotic cells for H3K9ac was highly significant (p-value < 10^−23^; two sided t-test).

**Figure S4. High concordance between mitotic and interphase histone modification patterns (biological replicate). (A)** Integrative genomic viewer (IGV) tracks showing the alignment for the indicated histone modification ChIP-seq results for interphase (blue) and mitosis (red) samples on an approximately 315Kb region on chromosome 1. The scale of each track was adjusted to the total number of reads using the Normalize Coverage Data option in IGV (see methods). The ChromHMM annotation for each genomic region is shown below the plot using the same color code as in **C**. **(B)** Scatter plots showing the normalized read counts for all the regions enriched in each modification (see methods for the peak calling details) in mitosis versus interphase. **(C)** Bar plots showing the percentage of reads in peak regions per ChromHMM category.

**Figure S5. TSS centered heat maps and metagenes.** Heat maps showing the normalized and standardized read depth at 6Kb regions centered at TSSs for all histone modifications that overlapped TSSs, for data obtained from two biological replicates **(A and B)**. Metagene plots showing the average read depth are below the heat maps.

**Figure S6. Enhancer centered heat maps and metagenes.** Heat maps showing the normalized and standardized read depth at 6Kb regions centered at enhancers for all histone modifications that overlapped enhancers, for data obtained from two biological replicates **(A and B)**. Metagene plots showing the average read depth are below the heat maps.

**Figure S7. Insulator centered heat maps and metagenes.** Heat maps showing the normalized and standardized read depth at 6Kb regions centered at insulators for all histone modifications that overlapped insulators, for data obtained from two biological replicates **(A and B)**. Metagene plots showing the average read depth are below the heat maps.

**Figure S8. H3K27ac does not penetrate the NDR during mitosis in G1E murine erythroblast cells.** Heat map showing the normalized and standardized read depth at 6Kb regions centered at TSSs using data from a previous study on G1E murine erythroblast cells (Hsiung et al. 2016). Metagene plot showing the average read depth is beside the heat map.

**Figure S9. Characterization of the TSS associated NDRs in a replicate experiment. (A)** A heat map showing the normalized and standardized z-scores at 6Kb regions centered at the TSS of 5731 genes that are occupied by H3K4me3, H3K9ac, and H3K27ac. The regions were manually divided into four groups and the percentage of regions in each group is shown in parentheses. **(B)** Metagene plots showing the average occupancy of each modification in each group. **(C)** Boxplot showing the difference in the GC content at the TSS for groups 1-3. The difference between group 1 and both 2 and 3 is highly significant (p-value = 2.01e-103 and 5.32e-64; t-test, and effect size = 0.76 and 0.95, respectively).

**Figure S10. Group 1 enriched motifs.** Motifs found enriched in group 1 versus the other groups, using the Discriminative Regular Expression Motif Elicitation (DREME) (Bailey 2011).

**Figure S11. TSS centered transcription factor ChIP-seq ENCODE data.** Metagene plots showing HeLa-S3 ENCODE ChIP-seq data for Groups 1-3. Only factors with counts higher than 10 are shown. See **Table S2** for a comprehensive list of p-values for all between-group comparisons.

**Figure S12. TSS centered histone modification ChIP-seq ENCODE data.** Metagene plots showing all HeLa-S3 histone modification ENCODE ChIP-seq data for groups 1-3. See **Table S2** for a comprehensive list of p-values for all between-group comparisons.

**Figure S13. Enriched GO terms found for the genes in** ***Groups 1*** **and** ***2.*** Biological processes enriched in *Groups 1* and *2* based on GO analysis. The x-axis is the −log of the FDR q-value as calculated by the GOrilla online tool (Eden et al. 2009). Note that *Group 3* did not yield significant results.

**Figure S14. Nucleosome shifting and deacetylation may explain NDR disappearance (biological replicate). (A)** Delta metagene plots showing the average read depth of the indicated modifications after subtracting the interphase depths from the mitotic depths. **(B)** A Scatter plot showing the normalized read counts at H3K9ac peaks on cells treated with TSA in mitosis versus interphase. **(C)** Metagene plot for H3K9ac data after TSA treatment. **(D)** Delta metagene plot for H3K9ac data before (dotted line) and after (solid line) TSA treatment.

**Figure S15. TSA treatment affects nucleosome positioning in enhancers and insulators.** Metagene plots of H3K9ac data after TSA treatment and delta metagene plots of H3K9ac data before (dotted line) and after (solid line) TSA treatment for NDRs associated with enhancers **(A)** and insulators **(B)**.

**Figure S16. TSA does not affect H3K4me3 localization.** A scatter plot showing the normalized read counts at H3K4me3 peaks on cells treated with TSA in mitosis versus interphase.

**Figure S17. Evaluation of the sensitivity of H3K9ac and H3K27ac antibodies to neighboring serine phosphorylation.** Epitope specificity of the indicated antibodies targeting H3K9ac and H3K27ac was assessed using histone peptide microarrays. These results revealed that nearby phosphorylation masks the epitopes for H3K9ac and H3K27ac antibodies. Comprehensive array datasets for these antibodies are deposited on the Histone Antibody Specificity Database (www.histoneantibodies.com).

**Figure S18. Mitotic release.** Cell arrested by STC were released by changing the medium and the percentage of non-mitotic cells was measured by fluorescence microscopy (using DAPI staining). The experiments were done on both TSA treated (green) and untreated (blue) cells. Each experiment was repeated twice and average and the standard error (error bar) are shown. TSA treatment significantly (p-value = 0.03, Paired Wilcoxon test) reduces cell ability to exit mitosis.

